# Establishment of hairy root culture and its genetic transformation of *Stephania tetrandra* S. Moore for production of BIAs

**DOI:** 10.1101/2023.05.17.541101

**Authors:** Xiuhua Zhang, Junling Bu, Yujun Zhao, Qishuang Li, Xinyi Li, Ying Ma, Jinfu Tang, Jian Wang, Changjiangsheng Lai, Guanghong Cui, Juan Guo, Luqi Huang

**Affiliations:** State Key Laboratory of Dao-di Herbs, National Resource Center for Chinese Materia Medica, China Academy of Chinese Medical Sciences, No. 16 South Side Street, Dongzhimen, Beijing 100700, China; School of Traditional Chinese Pharmacy, China Pharmaceutical University, Nanjing 211198, China; School of Pharmacy, Chengdu University of Traditional Chinese Medicine, Chengdu 610075, China

**Keywords:** *Stephania tetrandra* S. Moore, Fangji, hairy root culture, genetic transformation, benzylisoquinoline alkaloid, tetrandrine

## Abstract

Metabolic engineering improvement of plants will play an essential role in future agriculture, but this largely depends on the establishment of genetic transformation. *Stephania tetrandra* S. Moore is a traditional Chinese medicine used for rheumatalgia that accumulates benzylisoquinoline alkaloids as its main active ingredients. Wild or farmed plants have remained the main source of these essential medicines, resulting in supply pressure due to the scarcity of wild plant resources and the slow growth rate in cultivation. Here, we constructed *Agrobacterium rhizogenes* (C_58_C_1_)-mediated hairy root culture and a co-transformation system in *S. tetrandra* to obtain a new source of bisbenzylisoquinoline alkaloids production. We show that the biomass of the hairy roots increased by 10-fold, and the content of tetrandrine reached 8.382 ± 0.160 mg/g DW after 50 days of cultivation. In addition, overexpression of (R, S)-norcoclaurine 6-O-methyltransferase (6OMT) gene or treatment of hairy roots with methyl jasmonate (MJ) increased protoberberine alkaloid content. This work provides a method of obtaining hairy roots and a genetic transformation system for *S. tetrandra*, not only broadening the access to *S. tetrandra* resources, but also laying a foundation for further elucidation of the biosynthesis of tetrandrine and related alkaloids.

## INTRODUCTION

Fangji, the succulent taproot of *Stephania tetrandra* S. Moore, has been in widespread use over thousands of years in traditional Chinese medicine to treat rheumatalgia, arthrodynia, dropsy, dysuria, and eczema ^[1]^. There are abundant benzylisoquinoline alkaloids, including aporphines, proberberines, monobenzylisoquinolines and bisbenzylisoquinolines, in *S. tetrandra* ^[2]^, among which tetrandrine and fangchinoline are mainly responsible for antimicrobial, anticancer/antiproliferative, immunomodulatory, antidiabetic and neuroprotective activities ^[1,3]^. Pharmacological studies have indicated that tetrandrine shows anticancer activity by inhibiting fibroblast proliferation resulting from the modulation of tumor necrosis factor and collagen gene expression ^[4-6]^. In addition, tetrandrine is the most potent small molecule against Ebola virus and performs well in the treatment of silicosis in China.

Tetrandrine is currently extracted from wild *S. tetrandra*, in part because chemical synthesis of such a complex molecule (e.g., the coupling of two 1-benzylisoquinoline monomers and strict configuration requirements) is not commercially competitive. However, the root growth of *S. tetrandra* is very slow, and it takes at least five years to harvest. A slow growth rate and excessive exploitation have led to a shortage of wild *S. tetrandra* materials. Cell culture or hairy roots provide alternatives for the extraction of tetrandrine and other BIAs in *S. tetrandra*. In particular, hairy roots with a rapid growth rate, high biological activity, high biochemical stability, and the advantages of genetic transformation and so on have been constructed for many medicinal plants. In addition, hairy roots could be engineered to increase the content of active ingredients by overexpressing key enzyme genes or silencing competitive pathways. For example, the accumulation of benzophenanthridine alkaloids in the hairy roots of *E. californica* could be accelerated more than 5-fold by overexpressing BBE compared to control roots ^[7]^, and the contents of thebaine, codeine and morphine in the hairy roots of *P. bracteatum* were also increased by overexpressing SalAT ^[8]^. Moreover, with the application of scale-up bioreactor cultures, hairy roots such as ginseng ^[9]^ and *Polygonum multiflorum* ^[10]^ are being increasingly used to provide materials for the pharmaceutical and cosmetic industries. However, this relies on biosynthetic analysis of the active compounds in medicinal plants.

The biosynthetic pathways of bisbenzylisoquinoline alkaloids have been extensively explored. The upstream biosynthesis pathway of benzylisoquinoline alkaloids has been clarified in different species ^[11]^. Dopamine and 4-HPAA are condensed by (S)-norcoclaurine synthase (NCS ^[12]^) to produce norcoclaurine. (R, S)-norcoclaurine 6-O-methyltransferase (6OMT ^[13,14]^) catalyzes norcoclaurine to produce coclaurine, which is subsequently converted to N-methylcoclaurine by (S)-coclaurine N-methyltransferase (CNMT ^[15]^). N-methylcoclaurine is the precursor of benzylisoquinoline, protoberberine, morphinan, and aporphine alkaloids ^[16]^. N-Methylcoclaurine yields protoberberine and aporphine alkaloids through a reticuline intermediate, or the benzyl moieties of two N-methylcoclaurine units are oxidatively coupled by cytochrome P450 berbamunine synthase (BsCYP80A1 ^[17]^) to form bisbenzylisoquinoline alkaloids. Functionally characterized genes enable metabolic engineering of plants and hairy roots.

Since the first transgenic crop (the Flavr-savr tomato) emerged in 1994 ^[18]^, genome sequencing, bioinformatics analysis and genetic engineering technology have developed rapidly and been widely used in the fields of crop breeding and improvement ^[18]^ and plant chassis production of PNPs ^[19,20]^. Genetic editing of plants has been receiving increasing attention. At present, these biotechnology strategies are gradually being applied to medicinal plants ^[21,22]^, suspension cell culture and hairy root culture ^[23]^. However, the transformation of plants is often a technical bottleneck that restricts their development, which makes progress in plant genetic transformation systems crucial.

To the best of our knowledge, as an important medicinal plant for BIA production, there are no studies on the hairy root culture system and genetic transformation system in *S. tetrandra*. Here, we constructed the hairy root system of *S. tetrandra* by cocultivating leaf explants with *Agrobacterium rhizogenes* C_58_C_1_. The quantitative results showed that the content of tetrandrine in hairy roots reached 8.382 ± 0.160 mg/g DW. In addition, we successfully established a genetic transformation system of hairy roots through the transfer of eGFP and CyOMT-7 (6OMT in *C. yanhusuo*), which has been reported to be a key enzyme involved in the biosynthesis of BIAs ^[24]^. The results showed that exogenous genes could be successfully integrated into the tetrandrine genome and expressed. In summary, we established a hairy root culture system and a genetic transformation system of *S. tetrandra*, which will provide material and methods for the production of BIAs.

## METERIAL AND METHODS

### Materials

#### Plant materials

Seeds of *S. tetrandra* (local name: Fangji) were obtained from Yichun (28°11’33.53” N, 114°51’16.128” E), China, and grown in a greenhouse at the National Resource Center for Chinese Materia Medica, China Academy of Chinese Medical Sciences, Beijing, China.

#### Agrobacterium strain and binary vectors

*Agrobacterium rhizogenes* strain C_58_C_1_, which was used to induce hairy roots in *S. tetrandra*, was frozen and stored at -80 °C with 50% glycerin.

The binary vector pCAMBIA1300 with eGFP was from ph. Tang Jinfu. Screening resistance of the vector utilized kanamycin for bacterial culture and hygromycin for plant culture. The coding sequence of CyOMT7 was controlled by a super promoter in the binary vector pCAMBIA1300. Binary vectors pCAMBIA1300-eGFP and pCAMBIA1300-Cy6OMT-3-eGFP were introduced into the *A. rhizogenes* strain by the freeze‒thaw method.

### *A. rhizogenes-*mediated transformation

#### Preparation of infection solution

The wild-type *A. rhizogenes* strain was cultured in LB liquid medium supplemented with 50 mg/l rifampicin with shaking (200 rpm) at 28 °C under dark conditions. The cell density was adjusted to an OD600 of approximately 0.6-1.0 with LB medium. Before infection, acetosyringone (AS) was added at a final concentration of 100 μM to increase the efficiency of transformation. Then, the bacterial solution was centrifuged for 7 min at 4000×g to collect the incubated cells, which were suspended at a final cell density of OD600 =0.6 in MS+AS (100 μM) liquid medium for plant inoculation.

#### Induction of hairy roots

After disinfecting with 75% ethanol for 45 s and 2.5% NaClO for 7 min and cleaning with sterile water 3 times, the *S. tetrandra* leaves were cut into small slices of approximately 1 cm^2^ using sterile scissors. The injured leaves were submerged in infection solution, shaking (100 rpm) for 10 min at 28 °C. The explants were dried with sterile tissue paper and then placed on MS+AS (100 μM) medium with 0.8% (w/v) agar for cocultivation under dark conditions for 2 days at 25 °C. After 2 days of cocultivation, the explants were cleaned 3-5 times with MS liquid medium containing 500 mg/L cefotaxime and dipped for 5 min in MS liquid medium to remove excess cefotaxime. The explants were dried with sterile tissue paper and then transferred to selection medium (hormone-free half-strength MS medium containing 400 mg/L cefotaxime) for hairy root induction. The cefotaxime concentration was halved every 15 days. Many hairy roots emerged within 3-4 weeks from the wound sites. In addition, the induction of transgenic hairy roots requires the addition of 2.5 mg/L hygromycin to the screening medium.

The sterilized and vigorous hairy roots in solid medium were selected and transferred to half-strength MS liquid medium for expansion. The culture was incubated at 25 °C in the dark with shaking at a speed of 120 rpm and subcultured once every 21 days.

#### Hairy root growth curves

A total of 0.5 g (fresh weight) of *S. tetrandra* hairy roots that grew vigorously in half-strength MS liquid medium was transferred to 50 ml of new half-strength MS liquid medium and incubated at 25 °C in the dark with shaking at a speed of 120 rpm for 50 days. The hairy roots were harvested at 0, 5, 10, 15, 20, 25, 30, 40, and 50 d post inoculation. After recording the fresh weight, a small amount of the material was frozen in liquid nitrogen and stored at 80 °C for further use, and the remaining material was freeze-dried to detect the compound contents after recording the dry weight. Biomass is expressed as fresh and dry weight (DW) per 50 ml. The growth curve of the hairy roots was drawn from 0 to 50 days. The experiment was repeated three times.

#### Identification of transgenic hairy roots

The fluorescence of the transgenic hairy roots was preliminarily detected using a portable excitation light source (LUYOR-3415RG, Shanghai) with filter sets for eGFP (485/500 nm) or scanning confocal microscopy with filter sets for eGFP.

The hairy root genome was analyzed by PCR (polymerase chain reaction) for the presence of Ri plasmid fragments. Genomic DNA from the nontransformed roots of field-grown plants (negative control group) and four transgenic hairy root lines was extracted using a DNA extraction kit (Mai5bio, China). *Agrobacterium rhizogenes* C_58_C_1_ bacterial solution was used as a positive control for the bacterial DNA fragments Rol B, Rol C, and Vir D, and the vector pCAMBIA1300-Cy6OMT-3-eGFP was used as a positive control for the eGFP and Cy6OMT-3 genes. PCR with 2×EasyTaq^®^ DNA SuperMix (TransGen, Beijing) was run in a Veriti™ 96-well gradient PCR apparatus (Applied Biosystems, USA). The primers and fragment lengths are listed in Table 1. The reaction products and a standard DNA marker were visualized after electrophoresis in 1.5% agarose gels and photographed using the gel documentation system (Shanghai).

**Table 1.**
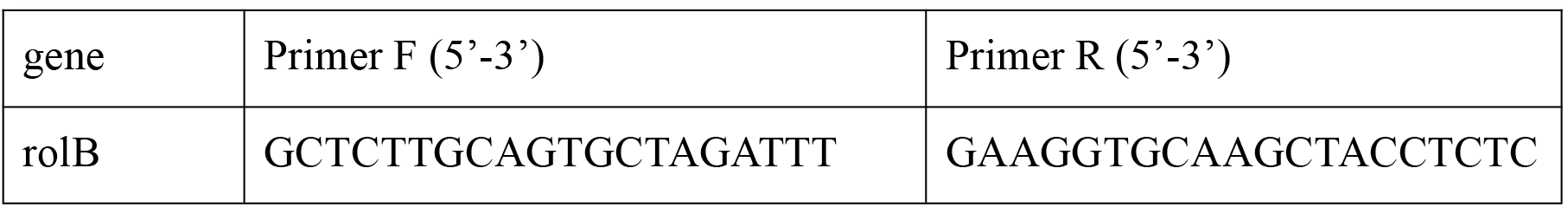
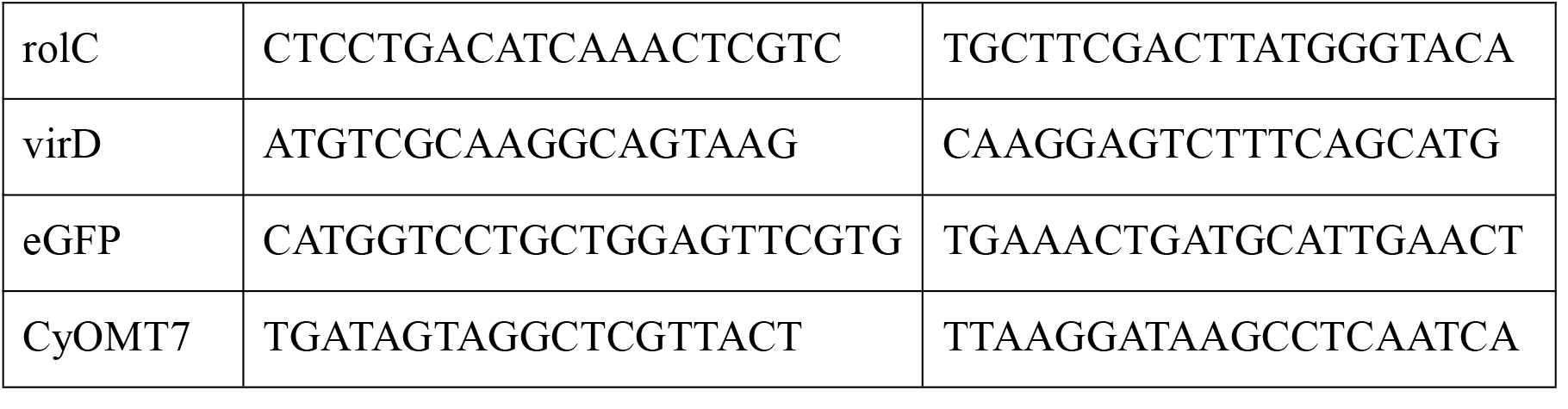
Primer sequences used for PCR analysis

### Hairy root treatment with elicitors

#### Elicitor preparation

Two elicitors, methyl jasmonate (MJ) and yeast extract (YE), were used in the elicitation process. The MJ solution (200 mM) was prepared in ethanol and filter-sterilized through a membrane filter (pore size: 0.22 μm). The YE stock solution (100 g/L) was prepared by dissolving YE in ddH_2_O at 121 °C over 15 min.

#### Elicitor treatment

One gram of 21-day-old hairy roots was cultured in 100 ml of half-strength MS liquid medium and incubated at 25 °C in the dark with shaking at a speed of 120 rpm for 14 days. MJ (50 μl; 0.1 mM final concentration) and YE (200 μl; 0.2 g/l final concentration) were separately applied to 14-day-old hairy root cultures. Equal amounts of ethanol and sterile water were added as controls. The hairy roots were harvested at 0, 1, 2, 3, 4, 5, 6, 8, and 10 posttreatment and washed with distilled water. The materials were frozen in liquid nitrogen and stored at 80 °C for further use. An aliquot of the material was freeze-dried to detect compound content. All experiments were performed in triplicate.

### Alkaloid extraction and quantitative analysis

#### Alkaloid extraction

Freeze-dried hairy root samples were crushed using a high-throughput tissue lapping device. Five milligrams of powder was accurately weighed, vortexed in 1 ml of 80% methanol and extracted by ultrasound for 30 min at room temperature; this procedure was repeated two times. Then, the samples were allowed to sit overnight. The extracts were separated by centrifugation and filtered through a 0.22 μm membrane filter prior to analysis.

#### Quantitative analysis

The extracts were quantitatively analyzed by LC-triple quadrupole MS. UPLC was carried out with an Acquity system using a CSH C18 column (2.1 mm × 100 mm, 1.7 μm particle size; Waters, Ireland). The mobile phases were acetonitrile (eluent A) and 0.5% aqueous formic acid (B) run at a flow rate of 0.4 ml/min with the following linear gradient elution program: 5% - 10% A from 0 to 3.0 min, 10% - 16% A from 3.0-5.0 min, 16% - 18% A from 5.0-18.0 min, 18% - 90% A from 18.0-23.0 min, 90% - 5% A from 23.0-28.0 min, and 5% - 5% A from 28.0-30.0 min. One microliter of sample was injected into the system. Sanguinarine (final concentration: 500 ng/ml) was added as an internal standard.

Eight target alkaloids [norcoclaurine (Yuanye, China), coclaurine (Yuanye, China), N-methycoclaurine (Rongchengxinde, China), fangchinoline (Yuanye, China), tetrandrine (Yuanye, China), reticuline (Yuanye, China), scoulerine (Rongchengxinde, China), and magnoflorine (Rongchengxinde, China)] were identified by the quantitative ion pairs 272.0→107.0, 286.0→107.1, 299.8→175.0, 609.2→367.2, 623,2→381.1, 329.8→192.1, 328.0→178.2, 342.0→297.1, respectively. Sanguinarine (500 ng/ml) was used as an internal standard with the quantitative ion pairs 332.2→274.1. Data acquisition and detection were performed in MRM mode. The data were processed using quantitative analysis software. For absolute quantification analysis of the target compound, the method was validated using a mixed standard solution, which was diluted with methanol to produce at least 5 data points.

### Statistical analysis

All experiments were conducted with at least three biological replicates. The BIA content was measured as the mean value ± standard deviation (SD). Error bars were determined for biological triplicates. The differences between the means were determined by analysis of variance with Tukey’s test using GraphPad Prism statistical software (version 7.0, USA).

## RESULTS

### Hairy root induction system in *S. tetrandra*

To obtain fast-growing materials for the efficient production of tetrandrine, we constructed a *S. tetrandra* hairy root induction system. In this study, the *A. rhizogenes* strain C_58_C_1_ was used to induce leaves of *S. tetrandra* to form hairy roots. The cut leaves of *S. tetrandra* were used as explants. Approximately 7-15 days after infection, calluses appeared at the wound sites, especially at the leaf veins (Fig. 2a, b). The calluses were white, solid and grew slowly (Fig. 2b). After subculture on cefotaxime-supplemented 1/2 MS medium for 20-30 days, approximately 1-2 hairy roots grew out on each callus (Fig. 2c). The *S. tetrandra* hairy roots showed typical morphological features with lateral branches and a lack of geotropism (Fig. 2d). The hairy roots identified as free of Agrobacterium contamination were transferred to 1/2 MS liquid medium (Fig. 2e), in which the hairy roots grew faster. In the process of culture, we found that the *S. tetrandra* hairy roots were relatively coarse and grew slowly, which is similar to the characteristics of the slowly growing, silty and swollen roots of *S. tetrandra*, suggesting that plant root traits may manifest in hairy roots.

**Fig. 1.**
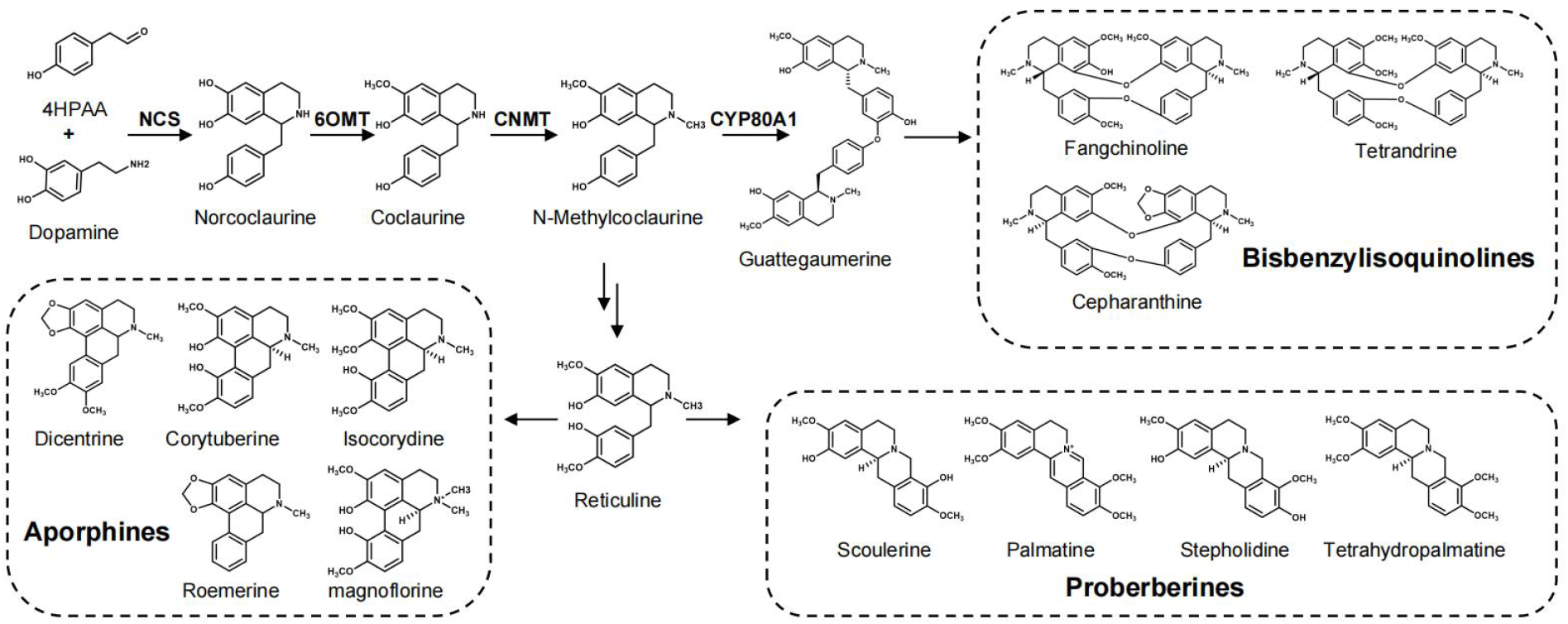
BIAs in *S. tetrandra* and their biosynthetic pathways.

**Fig. 2.**
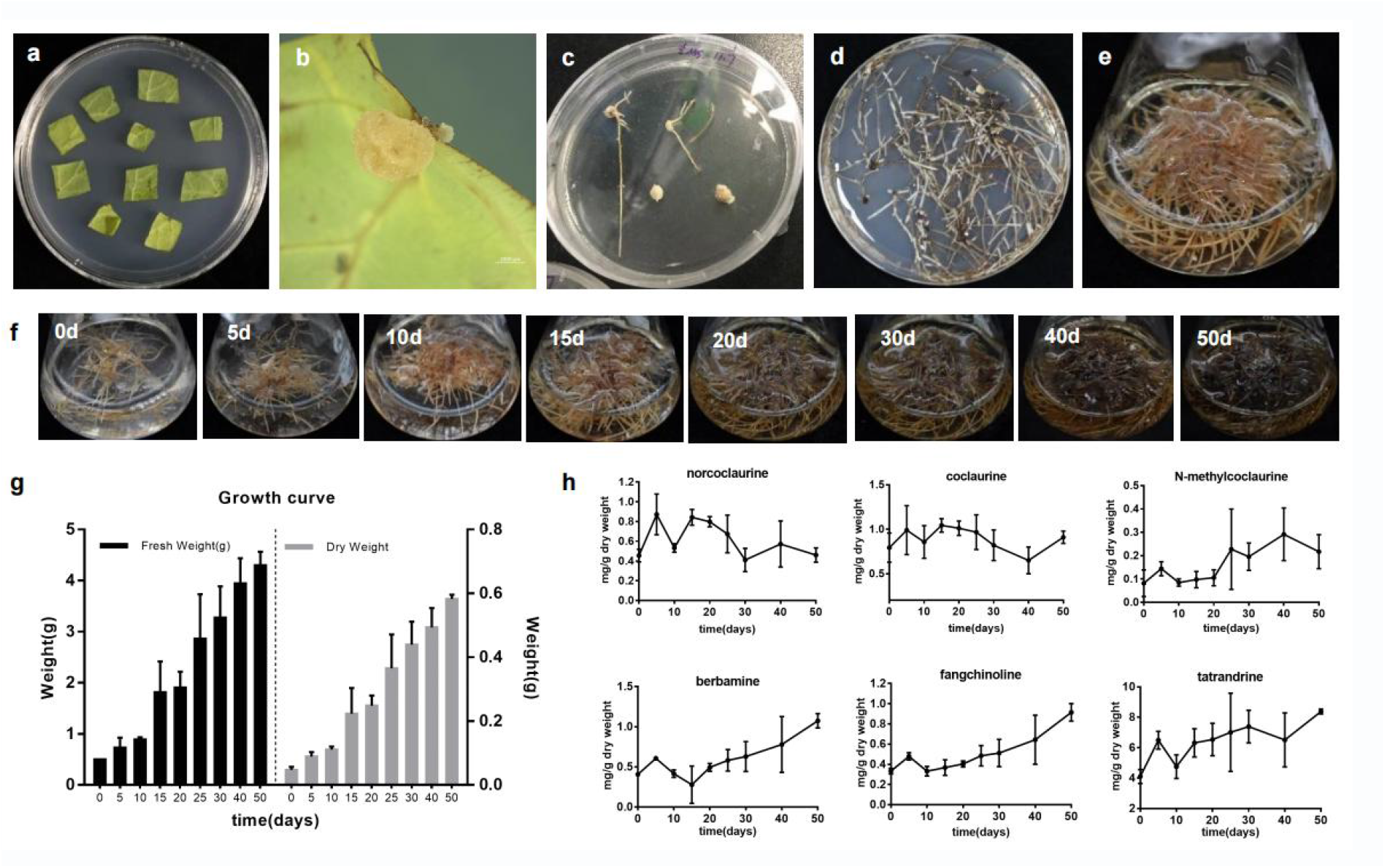
*A. rhizogenes*-mediated hairy root induction system in *S. tetrandra* and the consequent hairy root growth. Leaf explants (a), calluses (b) and hairy roots that appeared from the calluses (c). Hairy root culture on 1/2 MS solid medium (d) and in liquid medium (e). (f) Change in the growth status of hairy roots cultured in 1/2 MS liquid medium for 50 days. (g) Hairy root biomass growth curve. (h) Contents of alkaloids in the *S. tetrandra* hairy roots at different culture times. The scale bar on the figure is 1000 μm.

Hairy roots constantly exchange materials and energy with the external environment during growth. To explore the growth of hairy root culture, we investigated the biomass accumulation of the hairy roots after different suspension culture times in 1/2 MS liquid medium (Fig. 2f). Hairy roots that demonstrated stable repeatable growth were selected for continuous cultivation for 50 days, and their fresh and dry biomasses were measured. Without the addition of exogenous growth regulators to the medium, the growth of *S. tetrandra* the hairy roots exhibited a typical “S” curve (Fig. 2g). The fresh hairy roots grew slowly in the first 10 days and then rapidly from 10 to 30 days owing to vigorous cell division and adequate nutrition. After 30 days, the growth gradually slowed down, and the material started to brown. Finally, 4.30 ± 0.26 g FW and 0.582 ± 0.013 g DW were obtained after 50 days of cultivation, which were approximately 10 times the initial biomass. However, the 50-day-old hairy roots were dark brown and lacked vitality and could no longer be used for subculture, so expansion is appropriate at 20 days when the growth rate is the fastest.

To further determine the optimal harvesting time of the hairy roots, the BIA content in the hairy roots was compared between different culture times. The content of each compound did not change significantly in the first 20 days, and then the contents of precursor compounds (norcoclaurine, coclaurine) gradually decreased, and the biaBIAs (berbamine, fangchinoline, tetrandrine) began to accumulate. Contrary to the hairy root growth conditions, the rapid accumulation of biaBIAs mainly occurred after 30 days, which may be due to the stress experienced by the hairy roots caused by nutrient and air shortage in the later period. Eventually, the contents of fangchinoline and tetrandrine in the hairy roots reached 0.915 ± 0.087 mg/g DW and 8.382 ± 0.160 mg/g DW accounting for 14.62% and 59.67% of the content in the plants, respectively (content in plants was the average of plant content from each region ^[25]^). Therefore, *S. tetrandra* hairy roots should be harvested after 50 days.

In general, *S. tetrandra* hairy root culture is a practical and effective method for tetrandrine production, but further increasing the content of tetrandrine by genetic modification of *S. tetrandra* hairy roots is still needed.

### Hairy root genetic transformation system in *S. tetrandra*

Metabolic engineering has been used to modify plants and hairy roots to provide sustainable active compounds, such as *Artemisia annua* ^[21]^ and *Atropa belladonna* ^[22]^. However, genetic transformation of plants or hairy roots is needed. Therefore, we constructed a genetic transformation system for *S. tetrandra* hairy roots. The binary vector pCAMBIA1300 with an EGFP reporter gene was used for transformation, and the obtained hairy roots were examined by GFP fluorescence and genomic PCR. Compared with the wild-type hairy roots, the transgenic hairy roots showed bright fluorescence and contained EGFP gene fragments (Fig. 3a-e), which proved that *S. tetrandra* hairy roots had integrated the EGFP gene fragments carried by the overexpression vector from the engineered *A. rhizogenes*. In addition, the Rol B and Rol C genes, which are located in the T-DNA region of the Ri plasmid and were integrated into the plant genome to direct hair root differentiation, were present in the hairy roots, showing that hairy roots were induced by *A. rhizogenes* (Fig. 3e). The Vir D gene is required for T-DNA transfer and processing but is located outside of the T-DNA region and was absent in the transgenic hairy roots (Fig. 3e), showing that there was no *A. rhizogenes* contamination. In summary, the *A. rhizogenes*-mediated hairy root transformation system in *S. tetrandra* was successfully established.

**Fig. 3.**
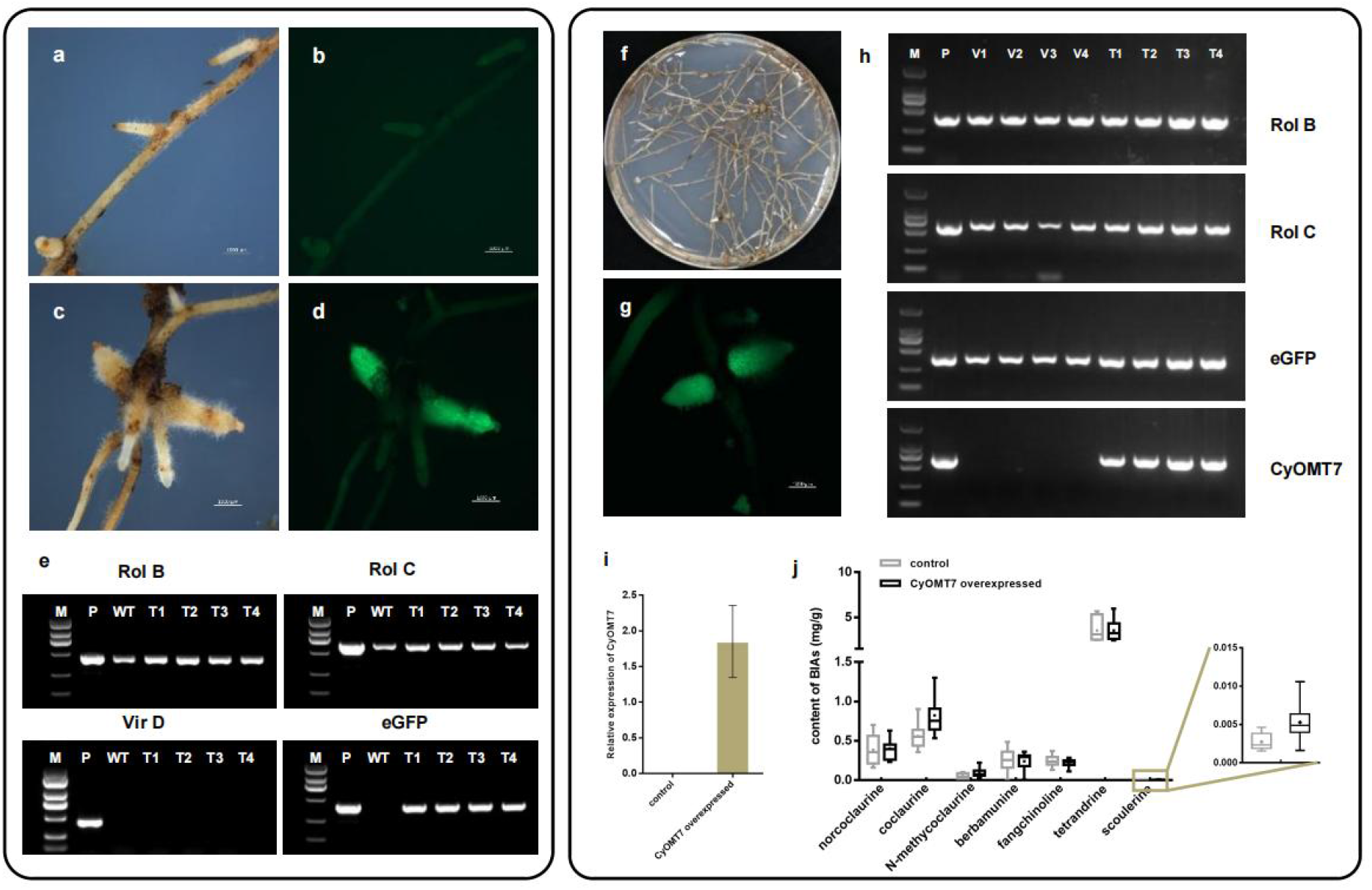
Establishment of the hairy root genetic transformation system. a-e: Hairy roots transformed by pCAMBIA1300-eGFP. WT hairy root (a) and GFP fluorescence (b). Transgenic hairy root (c) and GFP fluorescence (d). PCR analysis (e) of rol B, rolC, Vir D and eGFP in independent hairy roots transformed by pCAMBIA1300-EGFP. f-j: Identification of CyOMT-7-overexpressing hairy roots and determination of the compounds contained within. CyOMT-7-overexpressing hairy roots (f) and GFP fluorescence (g). PCR analysis (h) of rol B, rolC, eGFP and CyOMT-7 in CyOMT-7-overexpressing hairy roots. (i) Relative expression level of CyOMT-7 in control (hairy roots transformed by pCAMBIA1300-EGFP) and CyOMT-7-overexpressing hairy roots. (j) Contents of alkaloids in control and CyOMT-7-overexpressing hairy roots. The scale bar on the figure is 1000 μm.

We further confirmed whether the hairy root transgenic system could achieve functional verification of the target genes and metabolic engineering improvement of the hairy roots. According to previous research, changes in the expression of the 6OMT gene, which catalyzes the conversion of norcoclaurine to coclaurine, will affect the content of BIAs, such as sanguinarine ^[26-28]^. Here, CyOMT-7, the 6OMT in *C. yanhusuo* ^[24]^, was selected to verify the characteristics of this gene in the metabolic flow of *S. tetrandra* hairy roots for the following reasons: 1) the catalytic efficiency of CyOMT-7 is higher than that of St6OMT from *S. tetrandra* ^[29]^; and 2) alignment analysis revealed that there CyOMT-7 and the OMTs from *S. tetrandra* share 78% identity, which could avoid the gene silencing effect of endogenous genes. Ten independent lines were established as CyOMT-7 transformants, as confirmed by EGFP fluorescence and genomic PCR (Fig. 3f-h). RT‒qPCR analysis determined that CyOMT-7 was significantly overexpressed relative to the control group inoculated with pCAMBIA1300-EGFP (Fig. 3i). Quantitative analysis showed that the content of 1-BIA compounds downstream of the 6OMT gene, including coclaurine, N-methycoclaurine and scoulerine, a protoberberine-type compound, was higher in CyOMT-7-overexpressing hairy roots than in the control, while bisBIAs, including berbamunine, fangchinoline and tetrandrine, accumulated at similar levels in the transformants and the wild-type (Fig. 3j). This indicates that overexpression of CyOMT-7 is more favorable to the metabolic flow of protoberberine alkaloid synthesis.

### Effect of elicitor treatment on *S. tetrandra* hairy roots

The biosynthesis of secondary metabolites is affected by many factors. Elicitors, as extracellular signaling compounds, can trigger or initiate defense responses in plant cells, causing secondary metabolites to accumulate or be produced ^[30]^. Currently, the chemical elicitor MJ and biological elicitor YE are the most often used and have led to successful increases the contents of sanguinarine, dihydrosanguinarine and thebaine in poppy suspension cells ^[31,32]^.

After 15 days of culture, when the hairy roots grew rapidly and the change in compound content was relatively smooth (Fig. 2g-h), the hairy roots were treated with 0.1 mM MJ or 0.2 g/L YE for 10 days. In our study, 0.1 mM MJ treatment significantly inhibited hairy root growth and caused root browning (Fig. 4a). The effect of MJ on BIA accumulation mainly occurred during the first three days. Compared with the control treatment, the addition of MJ significantly increased the contents of coclaurine, *N*-methycoclaurine, tetrandrine, reticuline and scoulerine in hairy roots, which increased by 2.93, 5.65, 0.85, 2.75, and 13.44 times (Fig. 4b), respectively. However, there was no effect on hairy root growth or the accumulation of BIAs after treatment with 0.2 g/L YE (Fig. Supplementary).

**Fig. 4.**
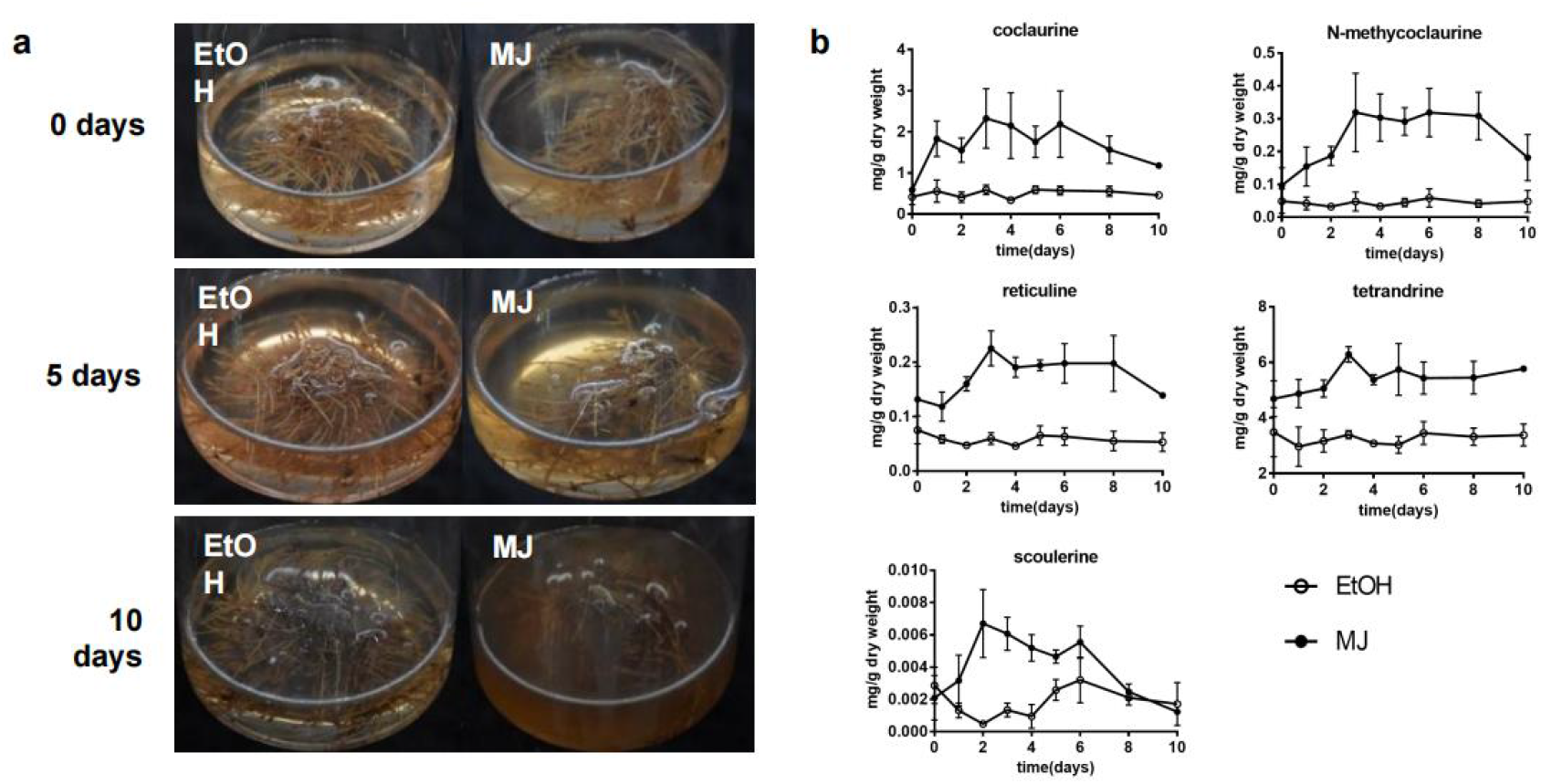
*S. tetrandra* hairy root culture elicited with MJ. (a) Hairy roots were treated with ethyl alcohol (EtOH, control) or methyl jasmonate (MJ) for 0, 5 and 10 days. (b) Contents of alkaloids in hairy roots treated with MJ for 10 days.

During tetrandrine biosynthesis, 0.1 mM MJ treatment significantly increased the contents of the precursor compounds (coclaurine, *N*-methycoclaurine). Although there was also an increase in tetrandrine, it was relatively weak. In a previous study, 0.1 mM MJ showed a weak increase in CYP80B1 expression ^[31]^, indicating that 0.1 mM MJ treatment may not be the most appropriate for tetrandrine production. The types and concentrations of the signal compounds and the leakage time are important factors that promote the production of secondary metabolites. Therefore, the conditions that can stimulate the synthesis of tetrandrine need to be explored. However, in our experiment and previous studies, it was found that 0.1 mM MJ significantly enhanced the content of protoberberines, which can be further applied to production.

## DISCUSSION

As a traditional Chinese medicine, *S. tetrandra* has great potential in the treatment of Ebola virus, silicosis and rheumatalgia. Due to its difficult cultivation, wild resources are heavily relied upon. However, this medicine is currently under great resource pressure, and an alternative material for production is urgently needed. In this study, we successfully established the *A. rhizogenes*-mediated hairy root transformation system in *S. tetrandra* (Fig. 5). The hairy roots grew rapidly, with a more than 10-fold increase after 50 days of culture. BIAs substantially accumulated in the hairy roots. Specifically, the content of tetrandrine in hairy roots was 8.382 ± 0.160 mg/g DW, which is slightly lower than that found in wild-type plant roots (this is an average of the contents in plants from each region ^[25]^). This shortage would certainly be overcome by the much higher growth rate of the hairy roots. Then, we established a hairy root genetic transformation system overexpressing CyOMT-7 (6OMT in *C. yanhusuo*), a key upstream enzyme gene, for further study of plant metabolic flow. Interestingly, the genetically modified hairy roots mainly increased the metabolic flow of protoberberine alkaloid synthesis rather than that of bisbenzylisoquinoline, similar to the MJ treatment experiment. This result is consistent with previous reports that overexpressing 6OMT or inducing *E. californica* suspension cultures with MJ resulted in an increased content of protoberberine alkaloids ^[28]^. These data indicate that the coupling and cyclization steps may be the rate-limiting steps that limit the synthesis of bisbenzylisoquinoline alkaloids in hairy roots and knocking down the branch pathway and overexpressing the BS gene may increase the tetrandrine content in hairy roots.

**Fig. 5.**
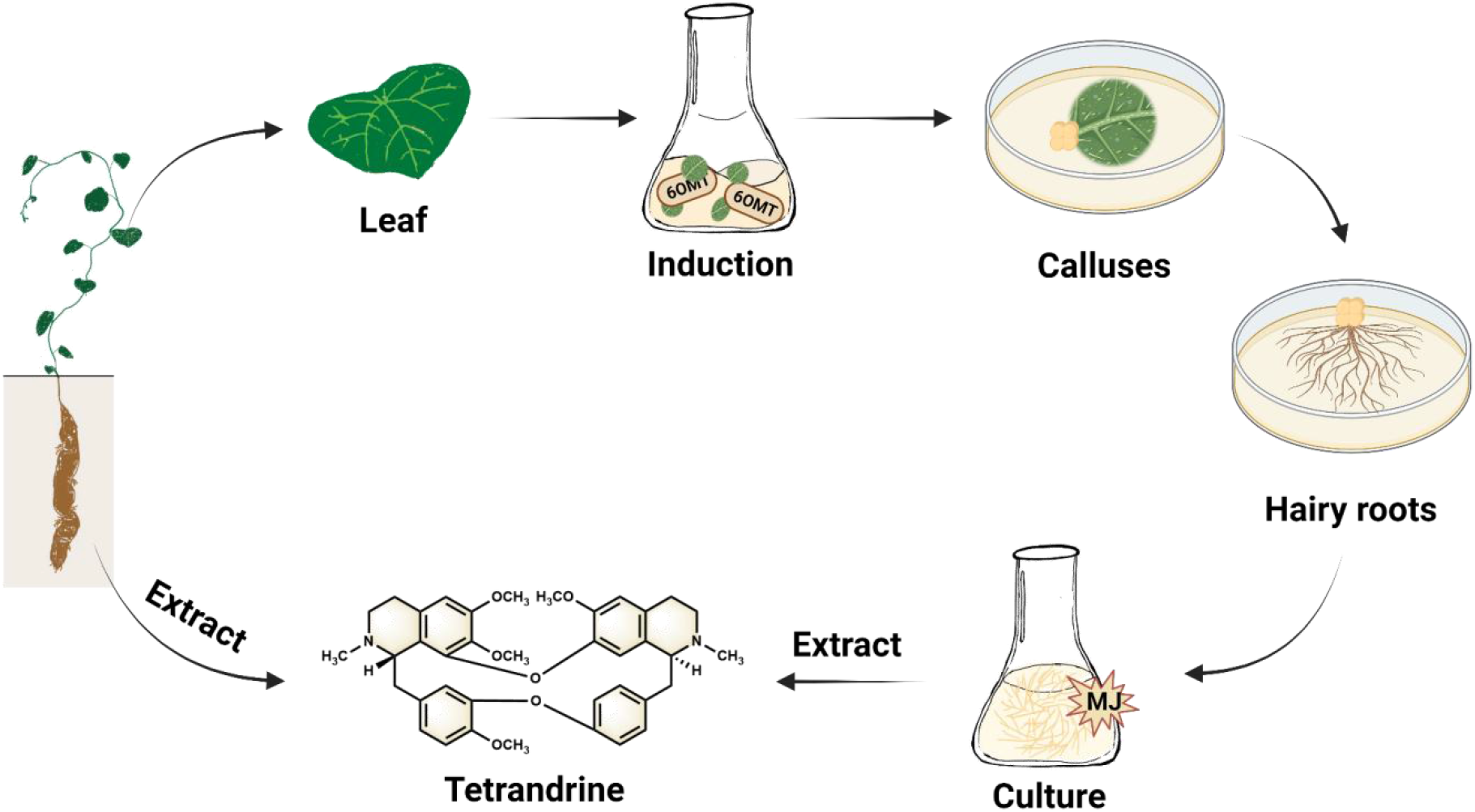
Hairy root culture and genetic transformation of *Stephania tetrandra* S. Moore

Active ingredients in medicinal plants are usually present in trace amounts and are tissue specific. With the development of plant tissue culture and genetic transformation technology, metabolic engineering has been widely used in the genetic modification of secondary metabolites in medicinal plants. For example, by simultaneously overexpressing multiple genes in the artemisinin biosynthetic pathway, HMGR, FPS and DBR2, Shen et al. obtained transgenic *A. annua* lines with significantly increased artemisinin contents ^[21]^. As a direction in plant metabolic engineering, hairy root genetic transformation systems have the advantages of a short transformation cycle, fast growth rate and high yield and have been applied to gene function characterization, secondary metabolite production, germplasm resource improvement and breeding ^[33]^, plant physiology and pathology research ^[34]^, showing great development value. At present, hairy root systems have been established in hundreds of medicinal plants ^[23]^, and many of them have been used in production. A variety of biotechnology strategies, such as multigene engineering, CRISPR/Cas9 and omics technologies, are gradually being applied to hairy root research, which will help us better understand and study important medicinal plants. In addition, in our study, we found that *S. tetrandra* hairy roots exhibited the same inflated, silty traits as the plant roots, implying that plant root traits may manifest in the hairy roots. Previously, we paid more attention to the secondary metabolism in hairy roots and used hairy roots to study the synthesis and metabolism of the active ingredients in medicinal plants ^[23]^, but this phenomenon indicated that hairy roots can be used to study the formation mechanism of root trait diversity in medicinal plants.

In conclusion, we have established the *S. tetrandra* hairy root induction system by coculturing leaf explants with *A. rhizogenes* C_58_C_1_ and further developed a hairy root genetic transformation system overexpressing the reported key enzyme 6OMT. The characteristics of a higher growth rate, effective transgenic methods and chemical compounds equivalent to those of plants not only address the challenges of *S. tetrandra* supply but also provide a method and technology system for research on *S. tetrandra*.

## Supporting information

Fig. Supplementary. S. tetrandra hairy root culture elicited with YE.

## Supplementary data

The following supplementary data are available at JXB online.

Fig. Supplementary. *S. tetrandra* hairy root culture elicited with YE.

## Acknowledgements

We thank all authors who kindly provided additional information and detailed data in their studies for this systematic review.

## Authors contribution

**Xiuhua Zhang:** Methodology, Investigation, Validation, Formal analysis, Data curation, Writing - original draft, preparation. **Junling Bu:** Methodology, Method guidance, Writing - review & editing. **Yujun Zhao and Qishuang Li:** Methodology, Method guidance. **Xinyi Li:** Methodology. **Ying Ma:** Writing - review & editing. **Jian Wang and Guanghong Cui:** Method guidance. **Jinfu Tang and Changjiangsheng Lai:** Resources **Juan Guo:** Supervision, Project administration, Conceptualization, Investigation, Validation, Results Interpretation, Writing - review & editing. **Luqi Huang:**funding acquisition, project administration, and resources. All authors contributed to the article and approved the submitted version.

## Data availability

All data generated or analysed during this study are included in this published article and its supplementary information files.

