## Supplementary material for "Establishment of hairy root culture and its genetic transformation of *Stephania tetrandra* S. Moore for production of BIAs": Fig. Supplementary. S. tetrandra hairy root culture elicited with YE.

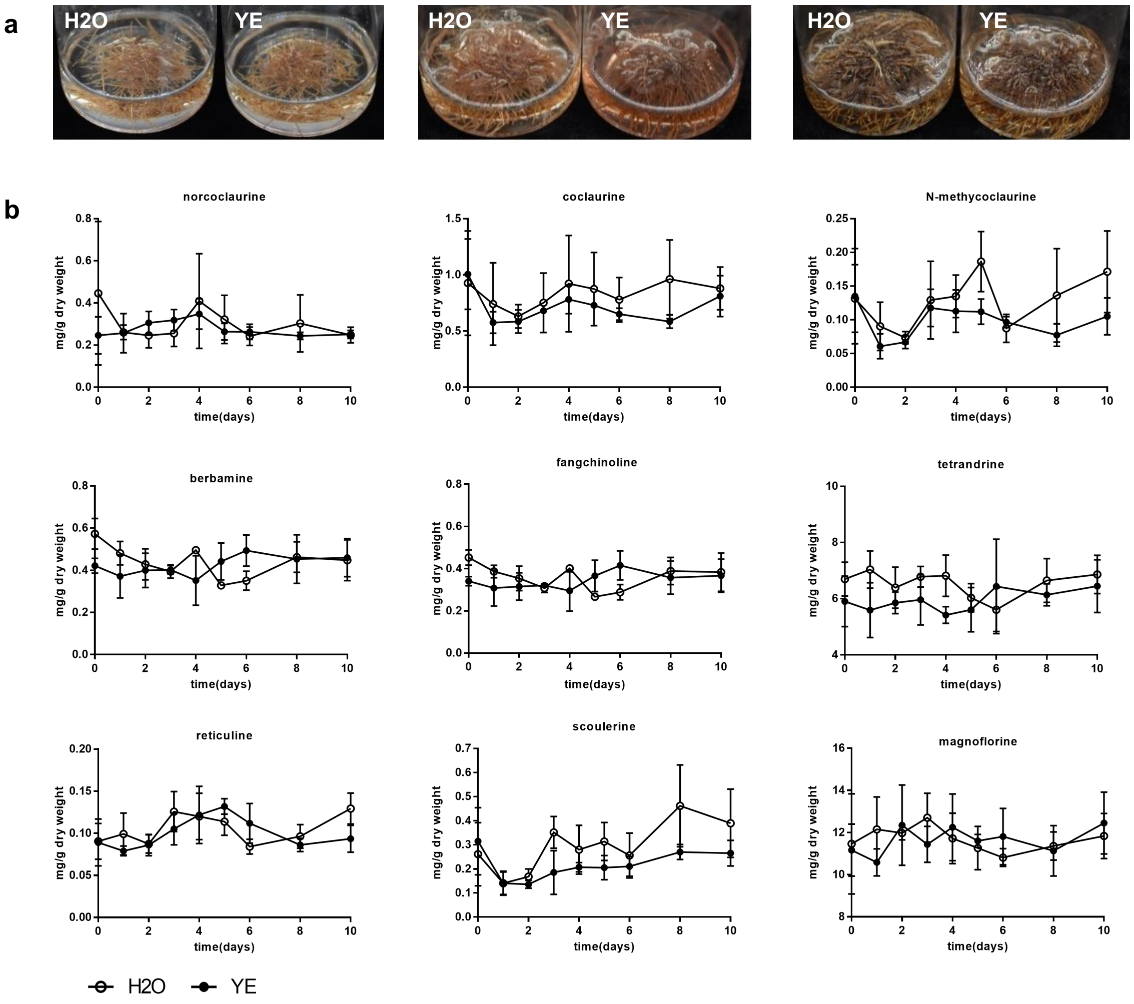


**Fig. Supplementary** *S. tetrandra* hairy root culture elicited with YE. (a) Hairy roots were treated with $H_2O$ (control) or yeast extract (YE) for 0, 5 and 10 days. (b) Contents of alkaloids in hairy roots treated with YE for 10 days.
